# Diagnosis-Optimized Dynamic Feature Learning Reveals Altered Default Mode Network Connectivity in Schizophrenia

**DOI:** 10.1101/2025.09.30.679557

**Authors:** Masoud Seraji, Charles A. Ellis, Liang Ma, Jessica A. Turner, Adrian Preda, Daniel H. Mathalon, Theo G.M. van Erp, Mohammad S.E. Sendi, Vince D. Calhoun

## Abstract

Schizophrenia (SZ) is characterized by widespread neural dysconnectivity, with particularly pronounced alterations in the default mode network (DMN)—a set of brain regions involved in self-referential thought and mind-wandering. We investigated dynamic functional connectivity within the DMN in SZ by analyzing resting-state fMRI data from two independent cohorts (FBIRN and COBRE). Using a novel iterative feature-removal clustering approach focusing on DMN independent components, we identified distinct recurring connectivity states while iteratively removing dominant connectivity features to reveal subtler network patterns. This approach iteratively refines the set of features considered, ensuring a more balanced representation and facilitating the identification of significant interactions that would otherwise be overlooked. Cluster centroids for the four connectivity states were highly similar across both datasets, reflecting stable shared DMN patterns. State occupancy was compared between participants with SZ and healthy controls, and associations with clinical symptom severity were examined. The SZ group exhibited significant alterations in DMN connectivity dynamics, spending more time than controls in certain connectivity states characterized by atypical DMN coupling and less time in a state reflecting a normative DMN configuration. A comparison between the FBIRN and COBRE datasets reveals both similarities and differences in occupancy rate (OCR) states. Overall, the pattern of group effects converged while state-specific divergences likely reflected protocol/sample differences rather than conflicting biology. Crucially, the explainability pipeline yielded a highly similar feature-removal order across datasets, indicating that the same DMN edges drive the diagnostic signal in both cohorts. Several OCR effects also tracked symptom ratings, linking abnormal DMN dynamics to clinical expression. Our findings suggest that SZ is characterized by reproducible disturbances in the temporal organization of DMN connectivity. These dynamic DMN features may serve as potential biomarkers for SZ, offering diagnostic and clinical utility by capturing network dysfunction that static connectivity measures could overlook.

## 1. Introduction

Schizophrenia (SZ) is a neuropsychiatric disorder affecting approximately 1% of the global population and is characterized by diverse clinical manifestations and widespread disruptions in brain dynamics ^1^. Schizophrenia presents a broad spectrum of clinical symptoms clustered into positive, negative, and cognitive or general domains ^2^. Positive symptoms include delusions, hallucinations, grandiosity, and increased suspiciousness or paranoia, among other manifestations ^3^. Negative symptoms are characterized by emotional withdrawal, blunted affect, reduced spontaneity or flow of conversation, and avolition. Cognitive deficits include attention, episodic memory retrieval, visuospatial processing, and decision-making deficits, as well as difficulties with abstract thinking ^4^. General symptoms often include increased anxiety, feelings of guilt, poor impulse control, and impaired judgment ^1^. These symptoms align with widespread neural disruptions, as SZ impacts multiple brain networks. Significant effects of SZ have been observed in various networks, including the visual, cognitive control, sensorimotor, auditory, and default mode networks (DMN). The DMN, in particular, has been a focus of study due to its strong association with self-referential thinking, memory processing, and mind-wandering. Alterations in DMN activity have been linked to deficits in cognitive processes and social cognition, which are common in SZ ^5^. Studies have highlighted the notable impacts of SZ on specific components of the DMN. For instance, there are marked disruptions in the anterior cingulate cortex (ACC), associated with emotional regulation and decision-making, leading to impairments in cognitive control ^6^. The posterior cingulate cortex (PCC) shows reduced connectivity that affects self-referential processing and attention ^7^. Additionally, the precuneus (PCN), crucial for episodic memory retrieval and visuospatial processing, exhibits altered activity and connectivity in individuals with SZ, contributing to deficits in these cognitive domains ^8^. Understanding disruptions within DMN subregions provides deeper insight into how their dysfunction contributes to the symptom manifestations of schizophrenia.

Neuroimaging modalities, particularly functional magnetic resonance imaging (fMRI), are crucial tools for investigating the neural underpinnings of SZ and other neuropsychiatric disorders ^9–13^. Both resting-state (rs) and task-based fMRI are employed for such analyses, but rs-fMRI is particularly valuable as it reflects the brain’s spontaneous activity patterns and is less influenced by task performance variability, which can differ significantly between participants with Schizophrenia (SZs) and healthy control participants (HCs), potentially confounding results ^14,15^. This modality’s ability to capture intrinsic brain activity makes it more suitable for studying neuropsychiatric conditions where task-based performance may be compromised ^16^. Furthermore, individuals with mental disorders are likely to perform tasks less effectively than HCs, introducing a potential confounder in any subsequent analyses ^17,18^. Several analytical techniques have been employed to process rs-fMRI data, notably independent component analysis (ICA) for identifying spatially independent brain networks and spectral analysis for examining the frequency-domain characteristics of brain activity ^19,20^. A prominent approach within rs-fMRI studies is functional network connectivity (FNC), which evaluates the co-activation and interactions between different brain regions or networks. FNC can be subdivided into static FNC (sFNC) and dynamic FNC (dFNC). sFNC assesses the overall correlation of network activity throughout an entire scan and has been applied to various disorders, including SZ, major depressive disorder, and Alzheimer’s disease, as well as to study cognitive processes like attention, executive function, and memory consolidation ^21–24^. However, dFNC, which tracks the fluctuation of connectivity patterns over time, offers deeper insights into the transient nature of brain network interactions and has proven effective in detecting disorder-specific changes that static analyses might miss [Allen et al., 2014; Damaraju et al., 2014; Fornito et al., 2015]. By capturing the evolution of transient connectivity between brain networks, dFNC can reveal subtle alterations in network interactions associated with disorders such as SZ [Sanfratello et al., 2019; Sendi et al., 2021a], providing a more nuanced understanding of brain function compared to sFNC ^27^. These dynamic insights are essential for uncovering the pathophysiology of complex disorders and informing future therapeutic strategies.

A common analytical approach for dFNC involves clustering techniques ^28,29^. These machine learning methods allow for the data-driven separation of dFNC data into clusters or states, with each dFNC time window assigned to a specific state ^11^. Individuals’ trajectories are quantified through these states using measures such as the occupancy rate (OCR)—the percentage of time spent in each state—and the number of state transitions, offering insights into the temporal dynamics of brain connectivity ^28^. However, clustering in dFNC analysis has inherent limitations. Certain features, especially those that show more pronounced variability, can disproportionately influence clustering results, potentially overshadowing significant but less variable network dynamics. This effect has been observed in whole-brain dFNC analyses where important SZ-related alterations in the DMN were overlooked but later detected in targeted DMN-specific studies ^7^. Additionally, findings from DMN-specific dFNC analyses have sometimes missed SZ-related effects that became evident only when less variable subregions were included in the study ^30^. This highlights the importance of methodological innovations that ensure a comprehensive analysis of all relevant network dynamics.

Previous studies have demonstrated that clustering results in dynamic FNC analyses can vary based on the brain networks or features included. However, such approaches may inadvertently obscure less dominant networks that could carry clinically meaningful, disorder-related signals ^31,32^. To address this, we introduce a novel feature learning-based approach in this study that emphasizes uncovering SZ-related dynamics within these less influential networks. Our method is adaptable for broader applications and involves an iterative process of clustering dynamic dFNC data, employing an algorithm-agnostic explainability technique to pinpoint the most influential feature affecting clustering outcomes. This top feature is subsequently removed, and the process is repeated until no further features remain for exclusion. Through this, we aim to provide a more comprehensive, network-wide analysis, capturing SZ’s full impact on brain connectivity (specially DMN) by integrating subtle network dynamics alongside dominant ones. Ultimately, our method demonstrates the potential to extract deeper, more widespread insights into how SZ affects brain function across networks, enhancing the understanding of its complex neural basis. By applying this approach, we aim to uncover nuanced differences in DMN subregion dynamics between SZs and HCs, as well as explore relationships between these dynamics and SZ symptom severity—insights that may be overlooked when analyzing the DMN as a whole. To test the robustness and reproducibility of our method, we apply it across two independent SZ rs-fMRI datasets. Applying this method to two independent datasets ensures results replicate across datasets and supports the generalizability of the approach. The iterative feature reduction and explainability analysis provide a more detailed understanding of how distinct subregions within the DMN contribute to SZ-related network alterations, potentially revealing previously obscured patterns critical for advancing research in neuropsychiatric disorders.

## 2. Method

In this section, we outline our study approach. First, we extracted dFNC data from SZs and HCs. Next, we assigned the samples to five clusters (states) and applied a feature learning method to iteratively remove the most important feature, followed by reclustering. This process continued until the number of remaining features made calculating the correlation distance impossible. After the feature learning phase, we computed OCR for each subject and state to examine potential changes in the temporal distribution of the five states across iterations. Finally, we compared the OCRs between the SZ and HC groups to identify significant differences and evaluated the relationships between the OCR values and symptom severity.

### 2.1. Study Participants

Data were obtained from the Mind Research Network Center of Biomedical Research Excellence (COBRE) ^33^ and the Functional Imaging Biomedical Informatics Research Network (FBIRN) ^34^. The COBRE dataset includes 89 HCs and 68 SZs. The FBIRN dataset contains 151 SZs and 160 HCs. The FBIRN raw imaging data were collected from seven sites: University of California, Irvine; University of California, Los Angeles; University of California, San Francisco; Duke University/University of North Carolina at Chapel Hill; the University of New Mexico; the University of Iowa; and the University of Minnesota. All participants provided written informed consent, and the institutional review boards at each site approved the consent process. In the FBIRN dataset, the diagnosis of schizophrenia was confirmed using the Structured Clinical Interview for DSM-IV-TR Axis I Disorders (SCID-IV) ^35^. Participants with schizophrenia were maintained on a stable dose of typical, atypical, or combination antipsychotic medications for at least two months prior to data collection, ensuring no significant changes in medication dosage during this period. Both SZs and HCs were excluded if they had a history of major medical illness, drug dependence within the last five years (excluding nicotine), current substance abuse disorder, or MRI contraindications. Additionally, SZs with significant tardive dyskinesia, HCs with a current or past history of major neurological or psychiatric illness, or those with a first-degree relative diagnosed with an Axis-I psychotic disorder were excluded. In addition, exclusion criteria for HCs included a current or past psychiatric disorder (except for one lifetime major depressive episode), head trauma with loss of consciousness greater than 5 minutes, recent substance abuse or dependence, depression or antidepressant use within the past six months, lifetime use of antidepressants for more than one year, and a history of a psychotic disorder in a first-degree relative. For the COBRE dataset, the inclusion criteria for patient selection required a diagnosis of schizophrenia or schizoaffective disorder in individuals aged 18 to 65 years. Each participant with SZ completed the Structured SCID-IV ^35^ with diagnostic confirmation achieved through consensus by two research psychiatrists. Patients had to demonstrate both retrospective and prospective clinical stability. Retrospective stability was confirmed by psychiatric records indicating no change in symptomatology or type/dose of psychotropic medications during the three months prior to referral. Prospective stability was assessed during three consecutive weekly visits and each imaging assessment. Exclusion criteria included a history of neurological disorders, head trauma with loss of consciousness exceeding 5 minutes, intellectual disability, or active substance dependence/abuse within the past year (excluding nicotine). Subjects with a history of dependence on phencyclidine, amphetamines, or cocaine, or use of these substances within the last 12 months, were excluded. A negative toxicology screen for drugs of abuse at the study’s onset was mandatory for all participants. HCs in the COBRE data set were recruited from the same geographic area through IRB-approved advertisements and completed the SCID-Non-Patient Edition to rule out Axis I conditions ^36^.

### 2.2 Data Collection

For the FBIRN dataset, imaging data was collected across seven sites. Six sites utilized 3-Tesla Siemens TIM Trio scanners, while one site employed a 3-Tesla GE MR750 scanner. The imaging protocol across all sites was standardized, using a T2*-weighted AC-PC aligned EPI sequence with the following specifications: TR = 2 s, TE = 30 ms, flip angle = 77°, slice gap = 1 mm, voxel size = 3.4 × 3.4 × 4 mm³, total number of frames = 162, and acquisition duration of 5 minutes and 38 seconds. These standardized parameters ensured consistency in data acquisition across multiple sites ^34^. The MRI images for the COBRE dataset were collected using a 3-Tesla Siemens Trio scanner equipped with a 12-channel radiofrequency coil. High-resolution T2*-weighted functional images were acquired using a gradient-echo echo-planar imaging sequence with the following parameters: echo time (TE) = 29 ms, repetition time (TR) = 2 s, flip angle = 75°, slice thickness = 3.5 mm, slice gap = 1.05 mm, field of view (FOV) = 240 mm, matrix size = 64, and voxel dimensions of 3.75 × 3.75 × 4.55 mm³. Resting-state scans included 149 volumes, during which participants were instructed to keep their eyes open and passively focus on a central cross (Aine et al., 2017). It is worth noting that while COBRE subjects were instructed to keep their eyes open during scanning, FBIRN subjects had their eyes closed.

### 2.3 Preprocessing

Statistical parametric mapping (SPM12, https://www.fil.ion.ucl.ac.uk/spm/) was used to preprocess the fMRI data ^20^. The first five dummy scans were discarded before preprocessing. Slice-timing correction was applied to the fMRI data, followed by rigid body motion correction to account for head movements in SPM. Next, the imaging data underwent spatial normalization to an EPI template in the standard Montreal Neurological Institute (MNI) space and was resampled to 3 × 3 × 3 mm³. Finally, a Gaussian kernel was applied to smooth the fMRI images with a full width at half maximum (FWHM) of 6 mm. To extract reliable DMN independent components (ICs) in each dataset, the NeuroMark automatic ICA pipeline within the group ICA of fMRI toolbox (GIFT, http://trendscenter.org/software/gift) was employed (Fig. 1, Step 2). This pipeline applies a spatially constrained ICA method using a previously validated NeuroMark_fMRI_1.0 template as a spatial prior ^37^. The template consists of robust and replicable group-level components derived from two large healthy control datasets. For each subject, the pipeline estimates individualized spatial maps and associated timecourses by fitting the group-level priors to the subject’s own fMRI data. From these subject-specific outputs, we focused on components that showed peak activation in gray matter regions consistent with the DMN. Based on an anatomical atlas ^38^, we further identified seven DMN subnodes of interest: three in the precuneus (PCN), two in the anterior cingulate cortex (ACC), and two in the posterior cingulate cortex (PCC).

**Fig. 1.**
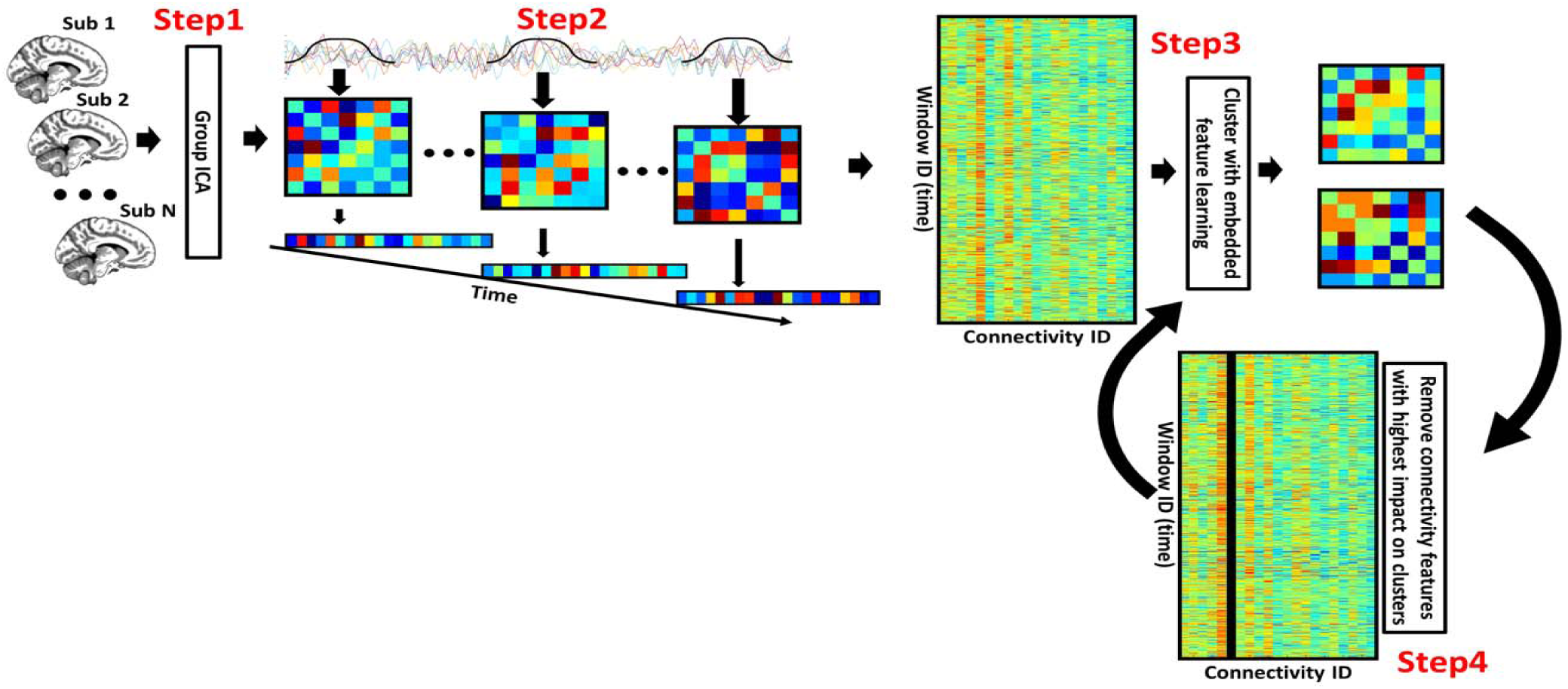
Overview of Methods. Step 1: After preprocessing, group independent component analysis (group-ICA) identifies default mode network (DMN) components and their time courses for all subjects. Step 2: Using a sliding-window approach, we compute dynamic functional network connectivity (dFNC)—pairwise correlations among DMN components—yielding a windows × edges matrix (rows = Window ID across all subjects/time; columns = Connectivity ID for each DMN edge). Step 3: Windows are clustered into connectivity. We then compute Global Permutation Percent Change (G2PC) by permuting one edge (or edge group) across windows, reassigning cluster labels with the trained model, and recording the percent of windows that change state; this ranks edges by global importance. Step 4: The top-ranked edge is ablated (removed/masked) and the data are reclustered, repeating Steps 3–4 until a compact, stable subset remains (e.g., until a pre-set number of edges or cumulative G2PC threshold is met).Note (black column): The black column in Step 4 marks the ablated connectivity feature (values set to zero and excluded from subsequent iterations), visually indicating its removal from the feature matrix prior to reclustering.

### 2.4 Dynamic Functional Network Connectivity Extraction

The dFNC of the seven subnodes in the DMN was estimated from each subject’s ICA timecourses using a sliding window approach, as illustrated in Figure 1. We employed a tapered window, created by convolving a rectangle (window size = 20 TRs or 40 s) with a Gaussian (σ = 3 s), to localize the data at each time point. The Pearson correlation was then calculated to measure the dFNC (Fig. 1, Step 2). The dFNC estimates from each window for each subject were concatenated to form a three-dimensional array (C × C × T), where C represents the number of subnodes (C = 7), and T represents the number of windows (T = 124 in COBRE and T = 137 in FBIRN). This array captured the temporal changes in brain connectivity between subnodes. Since the temporal resolution and eye condition differed between the two datasets, we analyzed them separately rather than combining them ^11,39,40^. This resulted in 21 connectivity features, with each feature representing the strength of the connection between pairs of DMN subnodes.

### 2.5 Feature Learning-based Clustering Analysis

After extracting the dFNC of the seven DMN subnodes, which resulted in 21 functional connectivity features, we applied k-means clustering with correlation distance to assign the samples into five clusters (Fig. 1, Step 3), as previously reported ^26^. To pinpoint the features most critical for clustering, we utilized the global permutation percent change (G2PC) feature importance method ^41^. This technique assesses cluster sensitivity by permuting each feature across all samples and measuring the percentage of samples that change clusters post-permutation (Fig. 1, Step 4). We then iteratively removed the highest-impact edges and reclustered, continuing until only two edges remained (fewer edges preclude our correlation metric). We used 500 random starts initially and, at subsequent iterations, initialized k-means from the previous centroids (excluding removed edges) to promote stability across iterations. Compared with a conventional single-pass k-means on all 21 edges, this feature-learning + reclustering framework (i) provides feature-level attribution of what drives each state, (ii) typically improves cluster separation and stability while preserving the canonical state topology reported in standard pipelines (See also ^41–43)^. Collectively, these steps enhance interpretability and robustness without sacrificing the reproducibility of dFNC states.

### 2.6 State-Related Analyses

Following the identification of network states using our feature learning approach, we aimed to compute the prevalence and consider the implications of these states in different subject groups. This section details our approach to analyzing state occupancy rates and their significance in distinguishing between SZs and HCs, as well as examining the relationship between these states and symptom severity.

#### 2.6.1 OCR Calculation

To quantify the prevalence of the identified network states, we calculated the OCR, which represents the percentage of time each subject spent in each state. This measure provided a basis for comparing how often SZs and HCs occupied each state, enabling us to investigate potential differences in network dynamics between the two groups. The OCR calculation was essential for subsequent analyses focused on disorder-related differences and symptom severity.

#### 2.6.2 SZ versus HC Analysis

We employed linear mixed models (LMM) to examine the differences in OCR between SZs and HCs for both the FBIRN and COBRE datasets. This approach allowed us to assess whether the diagnosis of schizophrenia was significantly associated with differences in state occupancy rates, while controlling for other variables. For the FBIRN dataset, the primary model used was OCR ∼ Diagnosis + Age + Sex + Site. In this model, ‘Diagnosis’ was the fixed effect of interest, while ‘Age’, ‘Sex’, and ‘Site’ were included as covariates. This setup enabled us to control for demographic variables and site-related variations, providing a robust comparison of state occupancy rates between SZs and HCs. The inclusion of multiple sites in the FBIRN dataset necessitated the inclusion of the ‘Site’ variable as a covariate to account for any site-specific effects. For the COBRE dataset, the model used was OCR ∼ Diagnosis+ Age + Sex. Since the COBRE dataset was collected at a single site, the ‘Site’ variable was not included as a covariate. This model focused on the effect of diagnosis on state occupancy rates while controlling for age and sex. By utilizing LMM for both datasets, we were able to account for the hierarchical structure of the data and provide robust statistical assessments of the observed differences in state occupancy rates between SZs and HCs.

#### 2.6.3 SZ Symptom Severity Analysis

To explore the relationship between symptom severity and OCR, we conducted separate analyses for the FBIRN and COBRE datasets. For the FBIRN dataset, we focused on the severity of both positive and negative symptoms. The models used were OCR ∼ Age + Sex + Site + PSS/NSS, where PSS (Positive Symptom Severity) or NSS (Negative Symptom Severity) served as the fixed effects of interest, with ‘Age’, ‘Sex’, and ‘Site’ as covariates. This approach allowed us to investigate the impact of symptom severity on OCR while accounting for demographic and site-related variations. By examining both positive and negative symptoms, we aimed to uncover how different aspects of symptomatology relate to state occupancy in SZs. For the COBRE dataset, we expanded the analysis to include general symptom severity (GSS) in addition to positive and negative symptoms. The models used were OCR ∼ Age + Sex + PSS/NSS/GSS, where PSS, NSS, or GSS were the desired fixed effects, and ‘Age’ and ‘Sex’ were the covariates. This comprehensive approach enabled us to explore the nuanced relationships between different dimensions of symptom severity and OCR.

## 3. Results

### 3.1 Cluster Centroids Analysis of the First Iteration in FBIRN and COBRE Datasets

As illustrated in Figure 2, the dFNC states identified in the FBIRN and COBRE datasets each reveal unique patterns of connectivity across various brain regions. These regions include the PCC, ACC, and PCN, highlighting distinct functional interactions within these networks. Notably, the first three states and the last state (State 1, State 2, State 3, and State 5) exhibit similar patterns across both datasets. This consistency suggests that these states reflect stable and common connectivity dynamics present in both populations.

**Fig. 2.**
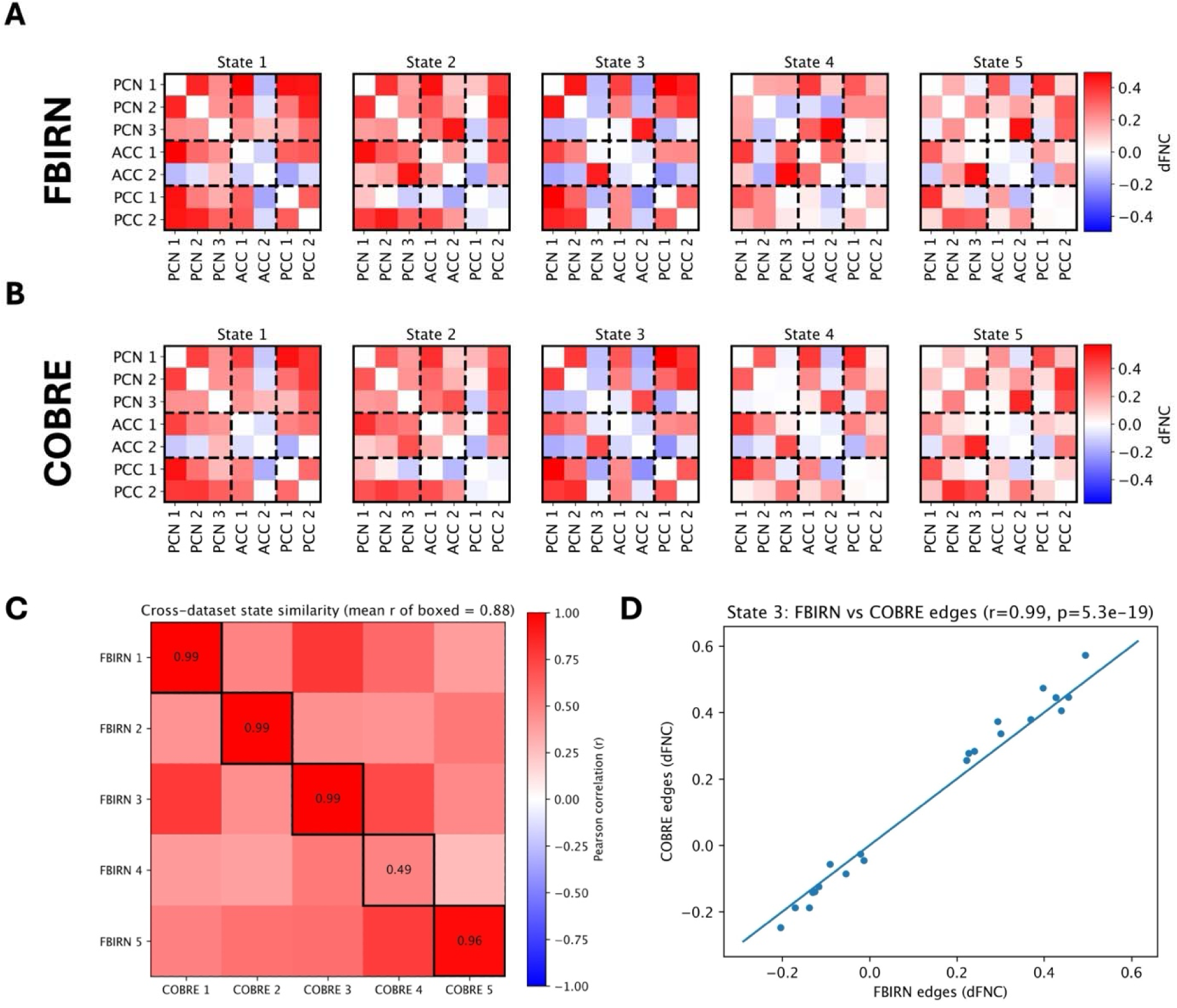
Cluster centroids of dFNC for the first iteration in FBIRN and COBRE datasets. Each heatmap represents the cluster centroids of pairwise dFNC values between different brain regions across five distinct states (State 1 to State 5) for both (A) The FBIRN and (B) COBRE datasets. The color intensity indicates the strength and direction of connectivity, with red representing positive connectivity, blue representing negative connectivity, and white indicating no connectivity. (C) Cross-dataset state similarity (Pearson r) computed from the vectorized upper-triangular edges of each centroid (rows = FBIRN, columns = COBRE). Black boxes mark the one-to-one matches; numbers in boxes are the matched correlations (mean r of boxed = 0.88). (D) Edge-wise correlation for the most similar pair (State 3): each point is one unique edge comparing FBIRN vs. reordered COBRE; line indicates identity. The association is very strong (r = 0.99, p = 5.3×10□^19^). The analysis reveals distinct patterns of connectivity for each state within both datasets, highlighting both similarities and differences in the functional network configurations.

In State 1, both datasets display strong positive connectivity within the PCN regions and moderate positive connectivity between the PCN and ACC regions, with the PCC regions exhibiting mixed connectivity with ACC and a positive connectivity with PCN. State 2 is characterized by a balance of positive and negative connectivity within the ACC region, while PCN regions maintain positive connectivity. State 3 highlights a mix of positive and negative connectivity within the PCN regions, with mostly strong positive connectivity between PCN and PCC regions, and weaker connectivity observed in ACC regions.

In both datasets, later-occurring states (States 4 and 5) were characterized by weaker and more fragmented patterns of connectivity compared to the earlier states. In FBIRN, State 4 exhibited relatively stronger coupling within the PCN, with reduced integration of the ACC and PCC, suggesting a posterior-dominant configuration. By contrast, COBRE’s State 4 was defined by globally weaker correlations across all DMN subregions, reflecting a low-cohesion or disengaged DMN state. State 5 displayed highly similar patterns in both FBIRN and COBRE. Across both cohorts, State 5 was marked by weak correlations among ACC, PCC, and PCN subnodes, with only faint residual coupling between posterior regions (PCN–PCC). This consistency across independent datasets suggests that a reproducible disengaged DMN states. The differences in States 4 suggest that while there are common connectivity patterns across both datasets, there are also population-specific variations in the dFNC states. The observed similarities in the other 4 states across both datasets highlight the stability of certain functional connectivity patterns. In contrast, the differences in the latter states underscore the influence of unique population characteristics on functional network dynamics.

### 3.2 Comparative Analysis of Occupancy Rates in FBIRN and COBRE Datasets

As we showed in Fig. 3, the OCR states observed in the FBIRN and COBRE datasets reveal significant differences between SZs and HCs (Blue representing negative t-values (HC < SZ) and red representing positive t-values (HC > SZ)). In the FBIRN dataset, five OCR states are identified. State 1 shows significant negative t-values, indicating that participants with SZ have a significantly higher occupancy rate in this state compared to HCs. States 2 and 5 show no significant differences, as noted in the absence of color (Except for the last iteration). States 3 and 4 reveal significant positive t-values, suggesting that HCs have a significantly higher occupancy rate in this state compared to participants with SZ.

**Fig. 3.**
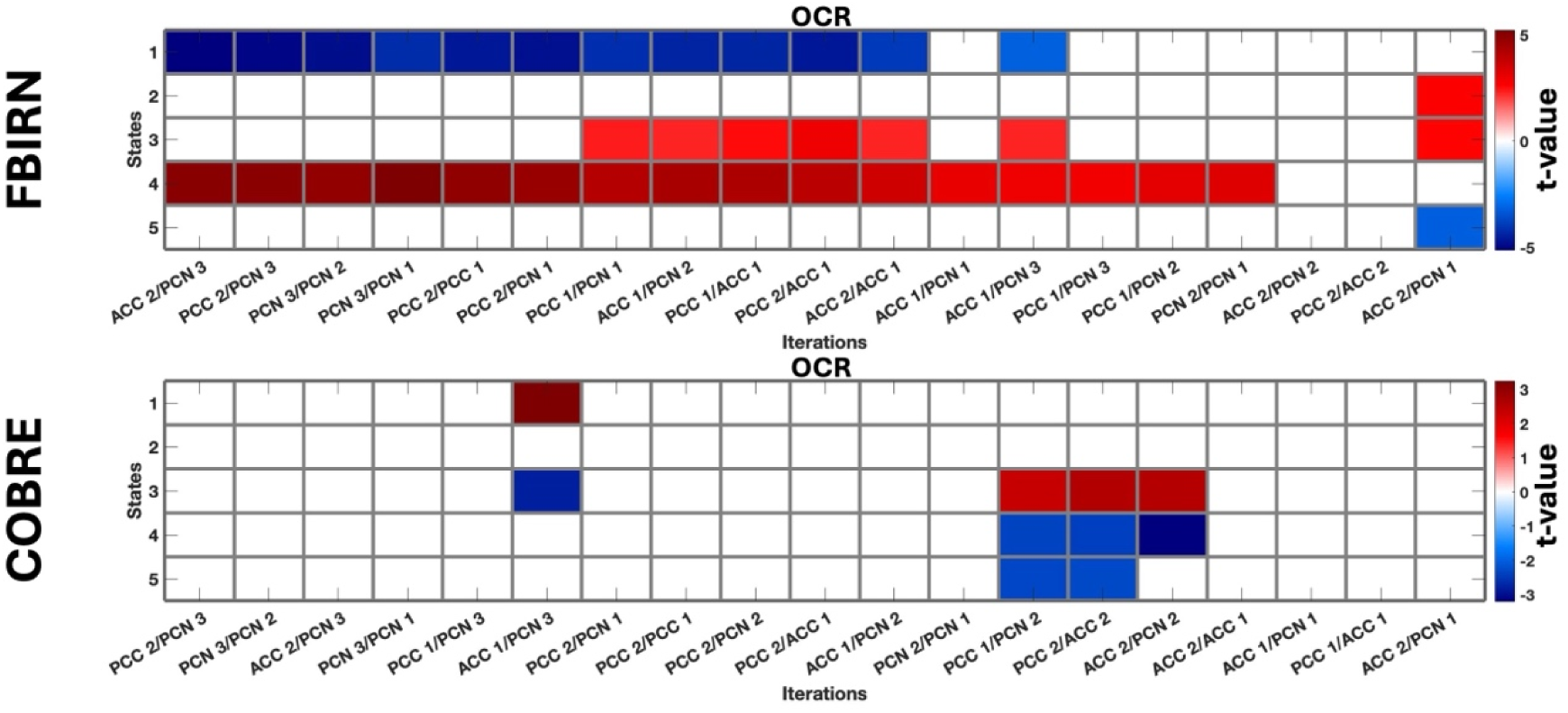
OCR States in FBIRN and COBRE Datasets. The figure shows the proportion of time subjects spent in specific functional connectivity patterns, with significant differences between SZs and HCs highlighted by blue and red colors. The colors indicate t-values, with blue representing negative t-values (HC < SZ) and red representing positive t-values (HC > SZ). Significant differences are observed after FDR correction. States without color indicate no significant differences between SZs and HCs. In both datasets, State 3 shows significantly positive t-statistics (HC > SZ), consistent with a higher occupancy rate in healthy controls.

The COBRE dataset also identifies five OCR states. State 1 shows a significant positive t-value, indicating that participants with SZ have a lower occupancy rate in this state compared to HCs. State 2 does not show significant differences, as indicated by the absence of color. State 3 has both positive and negative t-values, indicating a mix of significant positive and negative t-values, which suggests complex interactions and occupancy patterns in participants with SZ. States 4 and 5 with significant negative t-values, indicating a lower occupancy rate in HCs.

A comparison between the FBIRN and COBRE datasets reveals both similarities and differences in OCR states. Both datasets highlight State 3 with significant positive t-values, indicating a higher occupancy rate in HCs. However, while State 4 in the FBIRN dataset shows significant positive t-values (Higher occupancy rate in HCs), the COBRE dataset shows significant negative t-values for the same state (Higher occupancy rate in participants with SZ). Additionally, State 1 in the FBIRN dataset shows significant negative t-values, indicating a higher occupancy rate in participants with SZ. In contrast, a significant negative t-value was observed for just one iteration in the COBRE dataset for this state.

To evaluate the consistency of our explainability-driven feature importance across datasets, we compared the order of feature removal during iterative clustering between COBRE and FBIRN (Fig. 4). A Spearman rank-correlation test across the 19 features indicated a significant positive association between the two ranking sequences (p = 0.0034; permutation p = 0.0039, 10,000 randomizations), consistent with the near-diagonal structure of the heatmap. Features that were eliminated early in FBIRN tended also to be removed early in COBRE, while features retained until later iterations in FBIRN similarly persisted longer in COBRE. This diagonal structure in the heatmap underscores the robustness of our method across independent cohorts. Despite this overall agreement, some differences emerged — for instance, the feature ACC 2/PCN 1 was removed significantly earlier in FBIRN than in COBRE, suggesting it exerted a stronger influence on clustering in FBIRN. Conversely, PCC 2/PCN 1 was retained longer in FBIRN, highlighting dataset-specific variation in how certain DMN subregion interactions contribute to dynamic connectivity states. These nuanced differences may reflect subtle variations in population characteristics, scanning parameters, or network expression patterns across cohorts.

**Fig. 4.**
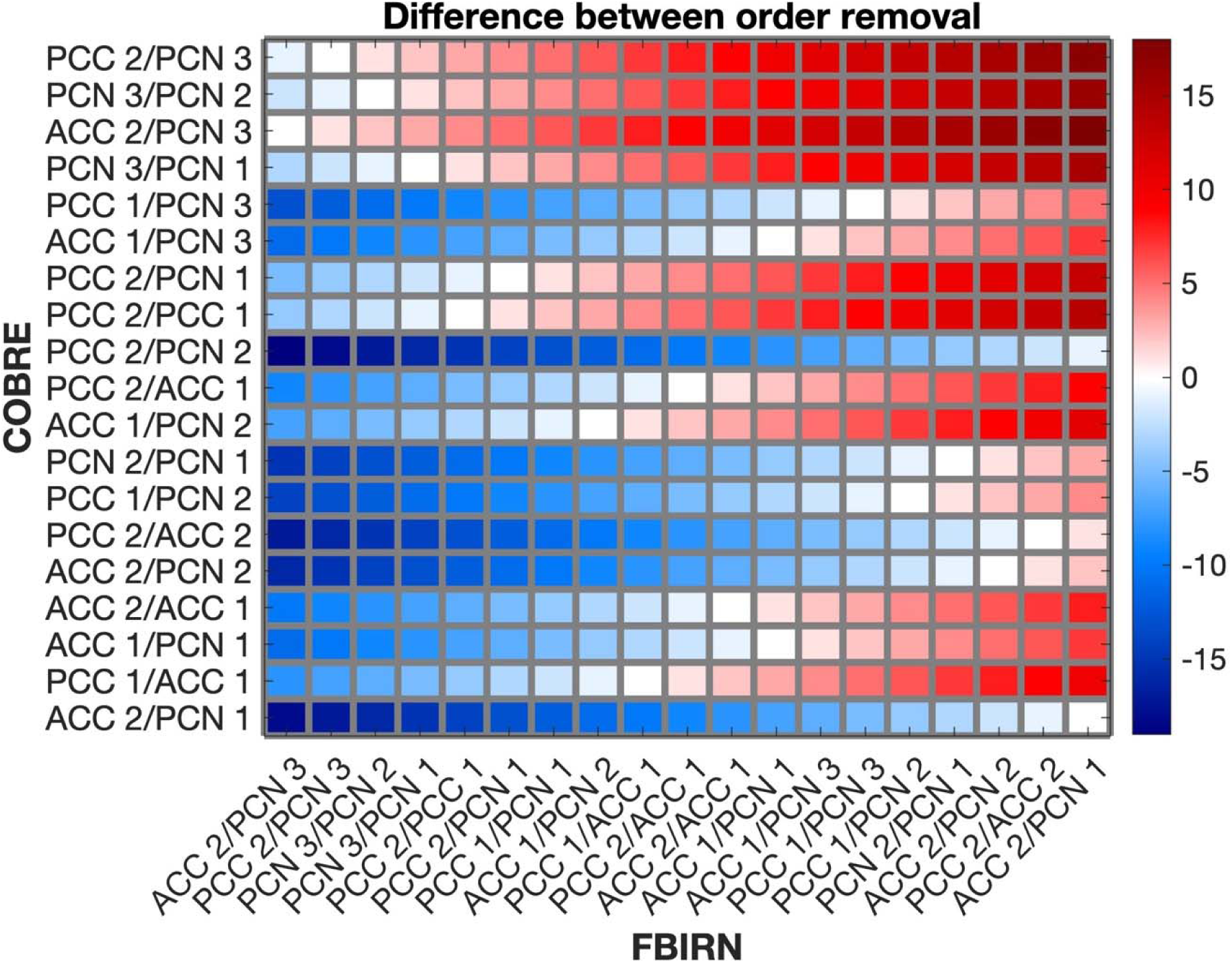
Heatmap of signed differences in feature removal order between COBRE and FBIRN datasets. Each cell quantifies the difference in the sequence of feature removal (importance ranking) between the COBRE (rows) and FBIRN (columns) datasets during iterative explainability-guided clustering. Positive values (red) indicate a feature was removed later in FBIRN than in COBRE; negative values (blue) indicate earlier removal in FBIRN. Global correspondence between rank orders was quantified with Spearman’s rank correlation (ρ = 0.64, p = 0.0034; permutation p = 0.0039, 10,000 randomizations). The diagonal trend from top-left to bottom-right suggests a strong correspondence in feature importance ordering between datasets, while off-diagonal deviations reflect dataset-specific effects.

### 3.3 Analysis of Symptom Severity in FBIRN and COBRE Datasets

To investigate the relationship between symptom severity and dynamic connectivity state expression in schizophrenia, we evaluated correlations between symptom scores and state occupancy rates (OCRs) across multiple iterations and connectivity features for both datasets. In the FBIRN dataset, we observed a limited number of significant associations. A late-removed feature, ACC 2/PCN 1, showed a positive association with positive symptom severity in State 5.

In contrast, PCN 3/PCN 1 and PCC 2/PCC 1 exhibited moderate associations with negative symptoms in States 2 and 3, respectively. In contrast, the COBRE dataset revealed broader and more statistically robust associations. For positive symptoms, both early- and late-removed features (e.g., PCC 2/PCC 1, ACC 2/ACC 1, ACC 2/PCN 1) were significantly associated with OCR across States 4 and 5, with some effects remaining significant after FDR correction (marked with asterisks). Notably, these results suggest that connectivity disruptions in both the anterior and posterior DMN subregions contribute to the expression of positive symptoms. Negative symptoms were also significantly associated with ACC 1/PCC 2 in State 2 and ACC 2/ACC 1 in State 5, highlighting distinct connectivity pathways. Interestingly, general symptoms in COBRE were primarily associated with reduced occupancy in State 1, driven by several early-removed features, including PCN-PCN and PCC-PCN interactions.

**Fig. 5.**
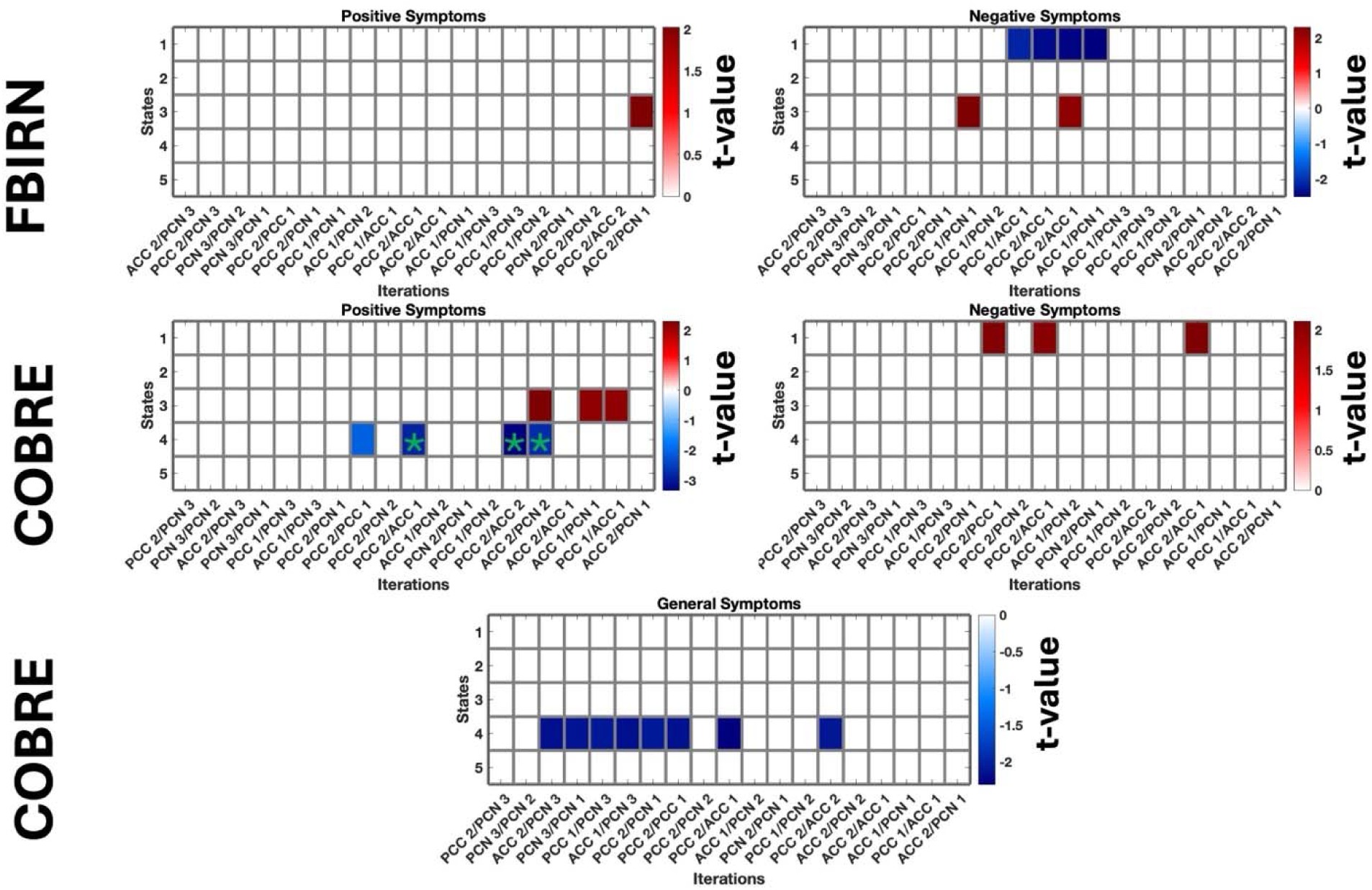
Relationship between symptom severity and dynamic connectivity state occupancy. Each subplot displays t-values from linear models relating symptom severity scores (Positive, Negative, and General) to occupancy rate (OCR) across clustering iterations and DMN connectivity features for FBIRN (top) and COBRE (bottom) datasets. Rows correspond to connectivity states, and columns denote removed features per iteration. Red and blue colors indicate positive and negative associations, respectively, with color intensity reflecting the strength of the t-value. Asterisks (*) mark features with statistically significant associations after FDR correction. While FBIRN shows sparse but localized associations, the COBRE dataset reveals broader and more statistically robust symptom-OCR relationships, particularly for positive and general symptoms.

Together, these findings indicate that symptom-related differences in network dynamics are more pronounced and widespread in the COBRE dataset, supporting the importance of considering dataset-specific variability when interpreting clinical relevance of connectivity features.

## 4. Discussion

In this study, we identified aberrant dynamic functional connectivity patterns within the DMN in schizophrenia. We demonstrated that these aberrations are reflected in the frequency with which participants with SZ occupy specific connectivity states. Using an explainability-guided, iterative clustering approach, we found that participants with SZ spent significantly more time in certain DMN connectivity states and less time in others, compared to healthy controls. Notably, across two independent datasets (FBIRN and COBRE), there was a convergent finding that one particular state (State 3 in our labeling) was under-occupied by SZ individuals (i.e. HCs dwell in this state more frequently). This state was characterized by relatively strong, well-balanced connectivity among DMN subregions – exceptionally robust PCN to PCC coupling – suggesting that participants with SZ have difficulty sustaining this “healthy” DMN configuration. Conversely, participants with SZ exhibited over-occupancy of other states, characterized by more atypical connectivity patterns. For example, in the FBIRN cohort, State 1 was occupied significantly more by SZ than HCs, whereas in COBRE, two states (States 4 and 5) showed greater occupancy in SZ. These differences suggest a redistribution of DMN activity in SZ, where patients spend more time in aberrant network states at the expense of engaging in more normative DMN configurations.

These findings resonate with and extend current models of SZ pathophysiology, particularly regarding the DMN’s role in the disorder. The DMN is a network implicated in self-referential thought, memory, and internally focused cognition, and it has long been observed to function abnormally in SZ ^26,44^. Prior static connectivity studies of SZ reported mixed results – for instance, some found hyperconnectivity within the DMN while others found hypoconnectivity, especially involving the ACC and PCC ^45,46^. Our dynamic approach helps reconcile such inconsistencies by showing that both patterns can occur in the same individuals, but at different times: participants with SZ alternate between states of heightened DMN coupling and states of diminished DMN coupling, with an imbalance in how long they can maintain the “healthy” versus “disconnected” states. In particular, our analysis of DMN subregions highlights an intriguing anterior–posterior imbalance in SZ. We observed that states dominated by strong posterior DMN connectivity (PCN–PCC coupling) with concomitantly weak anterior DMN (ACC) connectivity were often over-represented in SZ. This aligns with Sendi et al. (2021), who similarly reported that SZ is associated with increased connectivity between the PCN and PCC alongside decreased ACC connectivity ^26^. Furthermore, Sendi et al. validated their DMN findings across the same two datasets (FBIRN and COBRE), lending credence to the reliability of DMN dynamics as a marker of SZ. Our work adds that even when using a different analytic technique, one can achieve a high degree of replication (indeed, our identified state centroids were highly similar across datasets, with cross-site correlations ∼0.88–0.99 for matching states). This convergence across studies underscores a robust phenomenon: the DMN’s temporal coordination is consistently disrupted in SZ.

Our results align with and extend prior dFNC studies in SZ, and the comparisons yield important convergences and divergences. One clear convergence is that participants with SZ exhibit aberrant occupancy/dwell times across connectivity states, reflecting an altered brain network dynamics or “chronnectome” ^10^. Damaraju et al. (2014) analyzed whole-brain connectivity states and reported that participants with SZ spent *significantly less time* in certain high-connectivity global states relative to HCs. Those high-connectivity states in HCs were characterized by strong, cohesive interactions across multiple networks – essentially integrated brain states – which SZ individuals had difficulty sustaining. Instead, participants with SZ were more often found in states of dysconnectivity or fragmentary network coupling, and they transitioned more frequently into those aberrant states. Our DMN-focused findings echo this pattern: we also observed that HCs dwell longer in a state with robust DMN integration (State 3), whereas participants with SZ leave that state sooner and occupy other, less “normal” DMN states for a longer period. In fact, both datasets in our study identified a state that was preferentially expressed by HCs (with strong within-DMN connectivity) and a state (or states) preferentially expressed by SZ (with altered DMN coupling), paralleling the healthy vs. patient state dichotomy seen in whole-brain studies ^47,48^. Of course, not all studies find identical patterns, and our results also diverge in some respects from previous literature, reflecting nuances introduced by differences in analytical focus or sample composition. For instance, we found that one DMN state was more occupied by SZ in COBRE but less occupied by SZ in FBIRN (specifically, State 4 showed opposite group differences in the two cohorts). This initially appears contradictory but likely arises from subtle differences in how the states in each dataset were constituted (State 4 in FBIRN had a different connectivity signature than State 4 in COBRE; indeed, we noted these later states diverged between datasets). Such discrepancies highlight that dynamic states are not universal “atoms” of brain activity and can vary with analysis choices and cohort/scanner factors ^27,29,31^. Resting-state condition (eyes-open vs. eyes-closed) also modulates DMN connectivity and dFNC state expression ^49–51^. In addition, multi-site/hardware and protocol heterogeneity influence FC estimates ^52,53^, and illness stage/medication can further contribute to between-study differences ^5,15^. Despite these factors, both datasets here showed states with higher or lower occupancy in SZ, indicating consistent network dysregulation even when the exact state labels differ.

When broadening the perspective to other disorders, both parallels and distinctions emerge. Altered DMN dynamics are not exclusive to schizophrenia. For example, major depressive disorder (MDD) is also associated with abnormal DMN function – many participants with depression exhibit *hyperconnectivity* within the DMN and difficulty switching out of self-focused thought, which manifests as increased dwell time in DMN-dominated states (potentially underlying rumination) ^44,54,55^. However, the pattern in MDD appears to involve more uniformly elevated DMN activity, without the pronounced dissociation between the ACC and PCC observed. In bipolar disorder, dynamic connectivity studies have reported intermediate phenotypes: bipolar participants show disruptions in connectivity states, but often less severe than those in SZ and sometimes with unique features. One review noted that schizophrenia and bipolar disorder both exhibit altered dFNC, but SZ is marked more by large-scale network dysconnectivity, whereas bipolar disorder may show subtler dynamic “fluidity” changes ^56,57^.

In summary, comparing our results with prior dFNC studies reveals a coherent picture: schizophrenia consistently shows aberrant DMN connectivity dynamics, particularly an imbalance favoring internally focused, ACC-diminished states. This aligns with known DMN hyperactivity in psychosis ^44^ and reduced network integration in SZ ^10^. Our study adds a more detailed mapping of which DMN subconnections are most implicated and how these relate to symptoms. Divergences between studies (e.g., exact state definitions or whether SZ involves more or fewer state transitions) often stem from differences in methodology or scope, but they collectively emphasize that examining *when* and *how long* the brain occupies certain connectivity modes is crucial for understanding psychopathology. Our results encourage further cross-study integration – for instance, combining whole-brain approaches with focused network analysis – to piece together how DMN dynamics interplay with other networks in SZ. As the field moves toward increasingly large samples and multi-site consortium data, we expect that some of the subtle differences will resolve, and robust core findings (such as those we report for DMN) will stand as reproducible markers of the disorder.

### Clinical Implications

The demonstration of altered DMN dynamic connectivity in SZ has potential clinical implications, particularly for the development of biomarkers and novel intervention strategies. First, the fact that we could distinguish participants with SZ from HCs based on DMN state occupancy patterns – and do so in two independent datasets – suggests that these dynamic features hold diagnostic potential. In an era where psychiatry is seeking objective biomarkers to complement symptom-based diagnosis, measures like OCR of specific connectivity states could serve as candidates for identifying illness presence or quantifying severity. For instance, a patient who spends disproportionately little time in the “integrated DMN” state and excessive time in a “fragmented DMN” state might, on a neural level, be identified as having SZ (or at least a psychosis-spectrum condition) with a certain probability. This dovetails with recent machine learning efforts: models incorporating resting-state functional connectivity have successfully differentiated participants with SZ from controls with moderate accuracy, and dynamic features may improve that further ^58–63^. Our iterative feature learning method itself could be repurposed as a feature generator for classifiers. By ranking DMN connections by their disorder salience, it highlights which connectivity measures to feed into diagnostic algorithms.

Second, the associations we found between DMN dynamics and symptom severity hint at a role for these metrics in clinical monitoring and stratification ^26,64^. Because positive, negative, and general symptoms each showed relationships with different connectivity features or states, one could imagine using a panel of dynamic connectivity measures as an adjunct to symptom scales ^26,64^. For example, occupancy of a hyperconnected DMN state (with high PCC–PCN coupling) may serve as a biomarker for positive symptom intensity, possibly even detecting subclinical fluctuations in hallucination propensity ^65,66^. The link between negative symptom severity and ACC–PCC dysconnectivity suggests that patients who struggle to engage the anterior DMN may form a subtype that could benefit from targeted pro-cognitive or network-modulating treatments ^26,67^. Indeed, heterogeneity in SZ is a significant challenge; one patient’s illness can look very different from another’s. If validated, dynamic connectivity profiles could help parse this heterogeneity by providing neurophysiological signatures corresponding to different symptom dimensions (akin to how one might use inflammation markers in other fields to subtype a disease) ^64^. This approach aligns with the growing interest in neuroimaging-based patient stratification, where resting-state networks (including the DMN) are being explored for their prognostic value ^15,68,69^. For instance, Mehta et al. (2021) conducted a meta-analysis and found that dysfunction in the DMN (and striatal networks) was a consistent determinant of treatment response to antipsychotics ^15^. Our findings support this line of thought – if DMN dynamics reflect symptom severity, they may also reflect how a patient is responding to treatment (as symptoms ameliorate, one might expect partial normalization of DMN state occupancy). Further, this could be especially useful in guiding early intervention or recommendations for treatment-resistant cases: if a specific therapy increases the time the patient spends in a healthy DMN state, this change might be an early indicator of clinical response, even before symptoms improve. Conversely, persistence of abnormal DMN dynamics could flag non-response, prompting a change in therapeutic strategy.

Interestingly, our work also relates to the concept of transdiagnostic features. The DMN is a transdiagnostic network implicated in various psychiatric illnesses (schizophrenia, bipolar disorder, depression, PTSD, etc.), but how its dynamics manifest can differ ^5^. For example, hyperactivity of the DMN at rest has been proposed as an endophenotype for psychosis in general: it has been observed not only in chronic SZ, but also in individuals at clinical high risk for psychosis and in first-episode patients ^70–73^. This suggests that the tendency of the brain to fall into an overly active DMN state might predispose to psychotic experiences, regardless of formal diagnosis. Our finding that positive symptoms correlate with DMN state occupancy is in line with that idea, reinforcing DMN over-engagement as a candidate mechanism for psychosis risk across disorders ^65,66^. On the other hand, negative symptoms (and perhaps cognitive deficits) might align more with disorders like Alzheimer’s or major depression, where reduced activation of frontal networks and DMN hypoconnectivity are prevalent ^74–76^. It is notable that in our study, the general symptom factor – which encompasses cognitive and affective symptoms – was associated with reduced time in a strongly connected DMN state. This might connect with findings in dementia or severe depression, where the DMN’s integrity and activation capacity are diminished, correlating with global functional impairment.

It is also worth discussing the implications of our methodological innovation itself in a clinical context. The explainable feature-learning clustering gave us transparency into which connections are most disrupted in SZ. Interestingly, a number of the top features involved ACC–PCN or ACC–PCC links, which aligns with prior studies highlighting these connections as potential therapeutic targets or biomarkers ^26,65,77,78^. These could be followed up in future studies, measuring, for example, how those specific connections change with medication or predict longitudinal outcomes. If a particular DMN subconnection is reproducibly implicated, it might be measurable via electroencephalography (EEG or other modalities as well. For instance, the PCC–PCN coupling could relate to a known alpha rhythm coherence) ^79–85^. Thus, our work opens avenues for multimodal biomarker development: using the discovered features as hypotheses that can be tested with complementary techniques or simpler proxies ^86^. Finally, the replication across two datasets strengthens clinical utility. Reproducibility is a key barrier for translation; multi-site work shows that site/scanner factors can affect FC estimates, and robust biomarkers should generalize across such heterogeneity ^53,87^. Large-sample analyses further emphasize that reliable brain–behavior links require strong effects and adequate power ^88^, underscoring the value of findings that replicate across cohorts and pipelines.

### Limitation

Notwithstanding the strengths and implications discussed, our study has several limitations. First, there is the issue of dataset differences and generalizability. We purposefully analyzed two datasets (FBIRN and COBRE) to test reproducibility, and while many findings replicated, a few discrepancies arose. These discrepancies likely stem from differences in data acquisition and sample characteristics. For example, the COBRE scans were acquired with participants’ eyes open, whereas FBIRN used eyes closed. Eyes-open resting may lead to a slightly more externally oriented baseline, potentially affecting DMN activity levels (eyes-closed tends to increase DMN activation/spontaneous thought) ^49–51^. This difference could partly explain why certain state occupancies (like State 1 vs. State 4) showed opposite group effects between datasets – perhaps the eyes-open cohort had an easier engagement of a particular DMN state in HCs than the eyes-closed cohort did, or vice versa. Additionally, the FBIRN data were collected at multiple sites (seven scanning sites) with two scanner models involved, whereas the COBRE data were collected at a single site. We did include site as a covariate in FBIRN analyses, but multi-site data inherently have more noise and variance (scanner calibrations, environment, etc.) which could affect subtle findings. ^87,89^. Power also differed (FBIRN n = 151 SZ vs. COBRE n = 68 SZ), and smaller samples are more vulnerable to effect-size inflation and replication failures in brain–behavior associations ^88^. Thus, our most defensible conclusions are those that converge across cohorts, while dataset-specific divergences should be treated as hypothesis-generating.

A second limitation is related to the composition of the clinical sample. Both datasets consisted of chronic participants with SZ, most of whom were on antipsychotic medications at the time of scanning (maintained on stable doses) ^90,91^. Medication, especially dopamine antagonists, can modulate functional connectivity – some studies suggest antipsychotics normalize certain hyperconnectivities and alter network dynamics. Because all patients were medicated, we cannot disentangle which connectivity differences were due purely to illness and which might be influenced by treatment ^92^. Additionally, our samples did not include first-episode or unmedicated individuals; thus, we must be careful in extending conclusions to early-phase SZ or those who have never taken medication. Dynamic connectivity might look different in drug-naïve patients. A related point is comorbidity and other factors: we did not deeply characterize, for instance, cognitive functioning or substance use history in these analyses (though individuals with significant drug use disorders were likely excluded in the original studies). Cognitive deficits are a core part of SZ and correlate with functional connectivity measures; it would have been informative to examine how DMN state occupancy relates to cognitive test scores or daily functioning, but such data were not uniformly available. This represents an opportunity for future work but also a limitation in the present study’s scope ^93^.

Third, while our focus on the DMN allowed us to drill down into that network’s internals, it inherently provides a reduced field of view of brain connectivity. By restricting analysis to DMN ICs, we excluded other networks that are also implicated in SZ ^94^. It is possible that SZ has dynamic abnormalities in those networks or in the interactions between DMN and other networks (e.g., aberrant DMN–salience switching) that we did not capture here ^95–97^. We justified a DMN focus to reveal network-specific differences that whole-brain analyses might obscure, but this comes at the cost of not knowing whether a given DMN state co-occurs with a particular salience or executive configuration. Method choices in time-varying FC further influence which states and cross-network relations are detected, underscoring the need for complementary analyses^29,31^. In short, our study offers a detailed within-DMN picture but cannot determine how those states fit into the broader tapestry of multi-network dynamics and switching abnormalities in SZ; future work should integrate DMN with salience/executive systems in whole-brain dynamic frameworks ^10^.

Lastly, our symptom association findings, while intriguing, should be regarded as exploratory. The multiple comparisons problem is non-trivial – we tested correlations for three symptom dimensions across multiple states and features. We did use false discovery rate (FDR) correction and found several associations surviving ^98,99^, particularly in COBRE, but the FBIRN symptom results were sparse and did not strongly replicate COBRE’s pattern (possibly due to differences in symptom measures or sample size). Replication was also uneven across cohorts—likely reflecting limited power and measurement heterogeneity—consistent with broader concerns that robust brain–behavior associations and symptom links often require large samples and harmonized instruments ^87,88,100^. A further interpretational caveat is state dependence during “rest”: vigilance fluctuations and ongoing mentation (e.g., mind-wandering) can systematically modulate DMN connectivity during scanning, potentially biasing symptom–connectivity relationships ^101–103^. More broadly, causal interpretation is limited by our cross-sectional design and known statistical pitfalls in correlating symptoms with fMRI measures; longitudinal or interventional studies are needed to distinguish cause from consequence ^31,104,105^.

### Future Directions

Building on our findings and acknowledging the limitations, several avenues for future research emerge. A clear next step is to apply our diagnosis-optimized feature learning approach across broader conditions and populations. While we focused on SZ, the methodology is general and could be used to probe dynamic connectivity in other psychiatric or neurological disorders. For instance, applying this approach to bipolar disorder could elucidate which dynamic connectivity features distinguish bipolar from SZ – perhaps identifying shared vs. unique DMN alterations, which would inform the debate on the psychosis continuum ^57^. Similarly, in major depression or anxiety disorders, this method might uncover subtle network dynamics. By comparing feature-importance profiles across illnesses, we could determine if there are transdiagnostic dynamic biomarkers (e.g., DMN hyperconnectivity states in any condition with psychotic features) versus disorder-specific signatures. Ultimately, a comprehensive dynamic connectome atlas of different disorders could be constructed, guiding more precise diagnostics. In parallel, there is significant room to improve and expand the explainability and feature learning aspects of our approach. One direction is to incorporate more sophisticated explainable AI techniques. We used a straightforward algorithm-agnostic method to rank features. Future approaches could employ Shapley values or integrated gradient methods on a trained clustering model to assess feature contributions in a more nuanced way. Additionally, the iterative removal process could be refined – for example, instead of removing one feature at a time, one could remove or bin together highly correlated features or utilize domain knowledge (such as grouping edges by network pairs) to remove “features” in a cluster-wise manner. This might provide even more interpretable results.

Another future direction is moving towards real-time or longitudinal analyses. It would be enlightening to see if an individual’s state occupancy metrics change over the course of treatment or illness progression. A longitudinal study could scan patients at baseline, after initial treatment, and perhaps at follow-up, applying the same DMN state analysis. If, for example, effective treatment correlates with increased occupancy of the healthy DMN state, this would strongly support the clinical relevance of these metrics. Preliminary evidence in other contexts suggests that resting-state connectivity can change with therapy^15^. Our approach could detect even subtle shifts, given its sensitivity to small feature differences. Longitudinal data would also allow the construction of patient-specific trajectories: one could ask whether patients who remit (experiencing low symptoms) achieve a DMN dynamics profile closer to that of controls, whereas those with persistent symptoms remain locked in the pathological pattern. This could solidify the idea of using dynamic connectivity as a state marker (for current symptom status) and possibly as a trait marker if some abnormalities persist in remission.

We also advocate for future work to explore causal mechanisms underlying the observed dynamic patterns. One way is through multimodal studies: combining fMRI dynamic connectivity with EEG ^106–108^ or magnetoencephalography (MEG) could reveal the spectral or electrophysiological correlates of these states —e.g., DMN activity covaries with alpha/theta power and shows reproducible electrophysiological signatures across networks ^109–111^. For instance, MEG work has already examined dynamic network connectivity in schizophrenia at multiple timescales and in matched MEG–fMRI samples, demonstrating overlap and modality-specific differences ^25^. Extending such studies could test neurochemical links—for example, whether PET indices of dopamine (receptor availability or synthesis) track occupancy of particular connectivity states or modulate network coupling involving the DMN. Prior PET– fMRI work shows that dopaminergic manipulations and individual differences in dopamine relate to resting-state network connectivity, including DMN–control interactions ^112–114^. Understanding the biology could point to interventions: if, hypothetically, a certain oscillatory imbalance is key to sustaining an aberrant DMN state, neurofeedback or brain stimulation could target that oscillation ^82,115–117^.

Another future line is translating these findings to preclinical and animal models. While animals do not have a “DMN” in the human sense, rodents and primates do show analogous resting-state networks ^118–120^. It could be possible to induce a schizophrenia-like state in animals (via NMDA receptor antagonists, for instance) and use our dynamic analysis to see if their network dynamics shift in comparable ways (e.g., do ketamine-treated rodents show altered occupancy of a default-like network state?) ^121–124^. If so, this would create a platform to test new treatments and observe in real-time how they affect brain dynamics, thereby accelerating the bridge between bench and bedside for SZ therapeutics.

## 5. Conclusion

Our study demonstrates that schizophrenia is marked by aberrant DMN connectivity dynamics, evidenced by occupancy patterns that robustly differentiated patients from healthy controls in two independent cohorts. Critically, the clustering centroids were highly similar across four states in FBIRN and COBRE, indicating a stable, shared topology of DMN configurations. The iterative explainability pipeline produced a consistent feature-removal (importance) order across datasets, underscoring that the same DMN edges carry diagnostic signals despite differences in acquisition. Across cohorts, participants with SZ spent disproportionately more time in aberrant, less integrated DMN states and correspondingly less time in a normative, well-integrated state, with OCR showing both convergence and dataset-specific divergence (e.g., opposing effects in one later state), likely reflecting protocol and sampling differences rather than contradictory biology. State occupancy and salient DMN edges also tracked symptom dimensions—greater time in hyperconnected posterior DMN states aligned with positive symptoms, whereas reduced occupancy of the integrated state related to negative/general psychopathology—linking the temporal organization of the DMN to clinical expression. These reproducible state motifs, stable feature rankings, and symptom associations position dynamic DMN measures as promising biomarkers to augment diagnosis and monitor treatment response (e.g., normalization of time spent in the integrated state). Limitations include eyes-open vs. eyes-closed protocols, chronic medicated samples, and a DMN-only focus; nevertheless, the cross-dataset concordance of centroids and feature importance provides strong evidence that altered DMN dynamics are a reliable systems-level signature of schizophrenia.

